# P2X antagonists inhibit HIV-1 productive infection and inflammatory cytokines IL-10 and IL-1β in a human tonsil explant model

**DOI:** 10.1101/366179

**Authors:** Alexandra Y. Soare, Natasha D. Durham, Ramya Gopal, Benjamin Tweel, Kevin W. Hoffman, Julia A. Brown, Megan O’Brien, Nina Bhardwaj, Jean K. Lim, Benjamin K. Chen, Talia H. Swartz

## Abstract

HIV-1 causes a persistent infection of the immune system that is associated with chronic comorbidities. The mechanisms that underlie this inflammation are poorly understood. Emerging literature has implicated pro-inflammatory purinergic receptors and downstream signaling mediators in HIV-1 infection. This study probed whether inhibitors of purinergic receptors would reduce HIV-1 infection and HIV-1 stimulated inflammation. A human *ex vivo* human tonsil histo-culture infection model was developed to support HIV-1 productive infection and stimulated inflammatory cytokine interleukin-1 beta (IL-1β) and immunosuppressive cytokine, interleukin-10 (IL-10). This study tests whether inhibitors of purinergic receptors would reduce HIV-1 infection and HIV-1 stimulated inflammation. The purinergic P2X1 receptor antagonist, NF449, the purinergic P2X7 receptor antagonists, A438079, and azidothymidine (AZT) were tested in HIV-1 infected human tonsil explants to compare inhibition of HIV-1 infection and HIV-stimulated inflammatory cytokine production. All drugs limited HIV-1 productive infection but P2X-selective antagonists (NF449, and A438079) significantly lowered HIV-stimulated IL-10 and IL-1β. We further observed that P2X1- and P2X7-selective antagonists can act differentially as inhibitors of both HIV-1 infection and HIV-1-stimulated inflammation. Our findings highlight the differential effects of HIV-1 on inflammation in peripheral blood as compared to lymphoid tissue. For the first time, we demonstrate that P2X-selective antagonists act differentially as inhibitors of both HIV-1 infection and HIV-1-stimulated inflammation. Drugs that block these pathways can have independent inhibitory activities against HIV-1 infection and HIV-induced inflammation.

**Importance:** Patients who are chronically infected with HIV-1 experience sequelae related to chronic inflammation. The mechanisms of this inflammation have not been elucidated. Here we describe a class of drugs that target the P2X pro-inflammatory signaling receptors in a human tonsil explant model. This model highlights differences in HIV-1 stimulation of lymphoid tissue inflammation and peripheral blood. These drugs serve to both block HIV-1 infection and production of IL-10 and IL-1β in lymphoid tissue suggesting a novel approach to HIV-1 therapeutics in which both HIV-1 replication and inflammatory signaling are simultaneously targeted.

## Introduction

HIV-1 infection remains a major global health concern, despite the development of effective antiviral therapies to control the virus. An estimated 36.7 million people live with HIV-1, with 1.1 million people infected in the United States (1). Individuals on antiretroviral therapy (ART) can live long and healthy lives with suppressed viremia; however, infected individuals experience chronic inflammation associated with co-morbidities and increased risk of mortality (2–5). Despite undetectable viremia levels, long-term treated HIV-1 patients experience significantly higher rates of age-associated non-communicable co-morbidities (AANCCs), such as cardiovascular disease, frailty, and cognitive decline (6). The accelerated aging phenomenon has introduced new considerations in the care of HIV-1 infected patients (7–9).

The mechanisms underlying this chronic inflammation in HIV-1 infection are multifactorial. Depletion of CD4^+^ T cells at mucosal surfaces during HIV-1 infection can lead to reduced integrity of the mucosal epithelium and increased bacterial translocation (10). The subsequent elevated levels of plasma bacterial cell wall lipopolysaccharide (LPS) are associated with inflammatory biomarkers in HIV-1 patients, such as chronic monocyte activation, increased soluble CD14 (sCD14), and production of pro-inflammatory cytokines (11). Low-level viremia continues to stimulate systemic inflammation (12, 13). Despite numerous lines of evidence, no unifying mechanism has defined a connection between factors mediating early HIV-1 infection and cellular mechanisms of innate immune signaling.

Emerging literature has implicated the pro-inflammatory purinergic receptors in HIV-1 pathogenesis (19–35). Purinergic receptors mediate inflammation in many disease states (14–19) in response to extracellular nucleotides that are released from inflamed or dying cells (20, 21). P2X receptor subtypes are nonselective cation channels that can be found on a wide variety of tissue types, notably lymphocytes, monocyte/macrophages, and dendritic cells (DCs) (22–25). They are critical mediators of the innate immune response in a variety of different disease states including rheumatoid arthritis, transplant rejection, and inflammatory bowel disease (26–29). The P2X7 subtype is most highly expressed in immune cells (25, 30). P2X7 receptors, in concert with Toll-like receptors (TLRs), activate the NLRP3 (NACHT, LRR and PYD domains containing protein 3) inflammasome complex. The NLRP3 inflammasome is a highly conserved innate immune mechanism responsible for responding to pathogens by signaling cells to undergo pro-inflammatory cytokine production. This signaling mediates caspase-1-dependent release of IL-1β (23, 30), which can be secreted to promote inflammation or can stimulate pro-inflammatory lymphocyte programmed cell death known as pyroptosis which has been proposed to be an important cause of CD4^+^ T cell depletion (31, 32).

P2X1 and P2X7 subtypes are predominantly expressed on CD4^+^ T cells, the primary target of HIV-1 infection (42, 43). Recent studies by our group and others demonstrate that non-selective P2 antagonists blocking HIV-1 infection (33, 34). Non-selective P2X antagonists can reduce neurotoxic effects in murine neuron-microglial co-cultures exposed to HIV-1 transactivator of transcription (Tat) (35). These inhibitors can block HIV-1 infection in a dose-dependent manner during cell-to-cell and cell-free HIV-1 infection (34). Selective inhibitors of P2X receptors reduced HIV-1 replication in macrophages (36). Graziano *et al*. corroborated the importance of P2X7 in HIV-1 infection of macrophages by showing P2X7 inhibition blocked release of HIV-1 virions (37). Expression of P2X7 on human astrocytes is increased in the presence of HIV-1 Tat and P2X7 inhibitors have been demonstrated to reduce HIV-1 induced neuronal and microglial damage (38–40). Recently, Menkova-Garnier *et al*. demonstrated that P2X7 inhibitors restore T-cell differentiation in CD34^+^ cells derived from HIV-infected immunological non-responders (41). Additionally, P2X1 selective inhibitors were shown to inhibit HIV-1 fusion by blocking virus interactions with co-receptors C-C chemokine-receptor 5 (CCR5) and CXC chemokine-receptor 4 (CXCR4) (33, 42). As P2X receptors are also known mediators of inflammation and inflammatory signaling, it was of interest to understand whether HIV-1 stimulated inflammatory cytokine production would be abrogated by P2X inhibition.

Here, we examined the role of P2X-selective antagonists on HIV-1 productive infection and investigated whether these inhibitors block inflammatory cytokine production in response to HIV-1 stimulation. We demonstrate through an *ex vivo* tonsil model that these drugs can reduce HIV-1 stimulated IL-10 and IL-1β production, suggesting an important role for P2X inhibition in HIV-1 infection and HIV-stimulated inflammatory cytokine production.

## Results

### P2X inhibitors NF449 and A38079 can reduce HIV-1 productive infection in peripheral blood mononuclear cells (PBMCs)

Prior studies have reported that antagonists of proinflammatory purinergic receptor that detect extracellular ATP, here referred to as P2X inhibitors, can inhibit productive HIV-1 infection in T cell lines (33, 34, 43). We tested the role of a P2X1 inhibitor, NF449, and a P2X7 inhibitor, A438079, in blocking HIV-1 productive infection in PBMCs. Activated PBMCs were infected with HIV-1 NL-CI, a X4-tropic virus with an mCherry reporter, as previously described (44, 45), in the presence of NF449 and A438079. Reverse transcriptase azidothymidine (AZT) was tested as a positive control (Figure 1A). NF449 significantly reduced HIV-1 infection in PBMCs down to 25% while A438079 was less effective and reduced HIV-1 infection down to 60%. AZT inhibited infection by nearly 90%. None of the drugs tested exhibited toxicity on PBMCs (Figure 1B).

**Figure 1.**
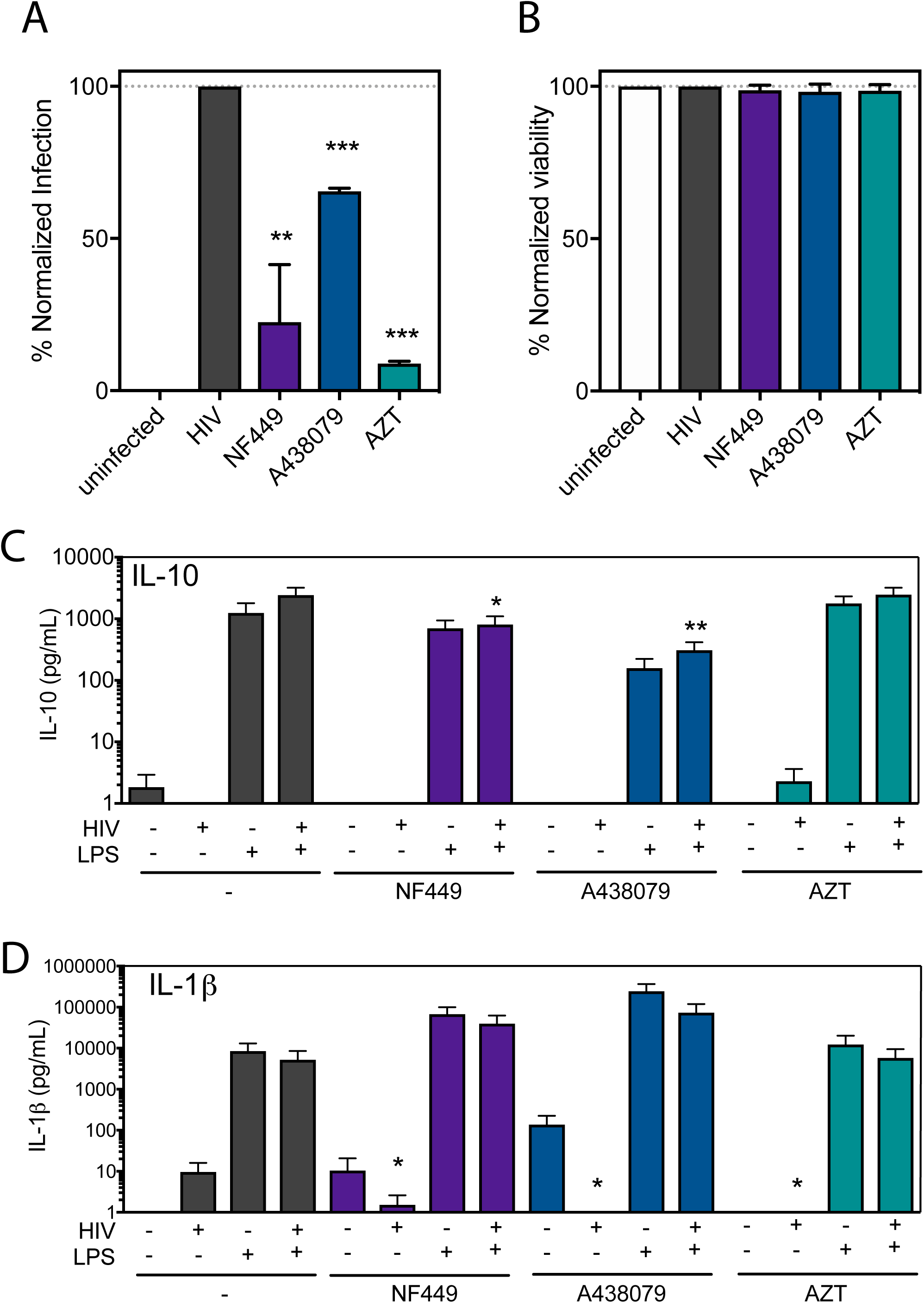
NF449 and A438079 inhibit HIV-1 productive infection in PBMCs with minimal inhibition of inflammatory cytokines. (A) PBMCs were isolated, activated, and infected with HIV-1 NL-CI (X4-tropic) for 48 hours. Cells were fixed and analyzed by flow cytometry then normalized to the HIV-1 infected condition for productive infection and (B) viability. (C) PBMCs were isolated and immediately exposed to HIV-1_MN_ (X4-tropic) for 12 hours in the presence or absence of LPS (1 pg/ml). Supernatants were collected and subjected to cytokine bead array (BD) for analysis of production of IL-10, IL-1b, IL-6, IL-8 IL-12, and TNF. Mean values ± SEM are represented for IL-10 and IL-1β from three donors. ^∗^, p ≤ 0.05, ^∗∗^, p ≤ 0.01, ^∗∗∗^, p ≤ 0.001.

Next, we tested the ability of these P2X-selective inhibitors to block HIV-1 stimulated inflammatory cytokine production. PBMCs were isolated and exposed to HIV-1_MN_ and tested for stimulation of inflammatory cytokines by cytokine bead array. Our initial findings indicated that minimal cytokine elevation was observed with HIV-1 infection of PBMCs (Figure 1C, 1D). As stated before, it is known that inflammasome activation requires two signals; the first is a TLR agonist and the second is a P2X agonist that we propose is stimulated by HIV-1 infection (46, 47). Therefore, we tested the effect of addition of a physiological level of LPS (1 pg/ml), a TLR4 agonist that has been reported to be circulating in the blood of HIV-infected individuals (10, 48, 49). Addition of LPS resulted in stimulation of IL-10 (Figure 1C) and IL-1β (Figure 1D) levels, but with only a small and not significant increase in HIV-1-dependent IL-10 production. We then pursued the establishment of a system more physiologically relevant to understand the interaction of HIV-1 infection and stimulation of inflammatory cytokine production.

### An *ex vivo* human lymphoid aggregate culture (HLACs) supports HIV-1 productive infection

*Ex vivo* infection of human lymphoid aggregate cultures (HLACs) with HIV-1 is a well-studied model system where HIV-1 induced-inflammasome activation has been characterized (50–55) and is an appropriate system to study inflammatory signaling that results from HIV-1 infection. Unlike blood-derived CD4-T cells, lymphoid-derived cells are not naturally resistant to pyroptosis in culture and do not require activation or addition of exogenous TLR agonists for HIV-1 infection (51). Human tonsil explants were obtained from healthy tonsillectomy patients, homogenized as previously described (56) and cultivated in HLACs. HLACs were infected with HIV-1 NL-CI and harvested on 0, 2, 5, 8, and 12 days post infection (DPI) (Figure 2A). Infection was quantified by flow cytometric detection of mCherry-positive viable cells as indicative of HIV-1 NL-CI infection (Figure 2B). Peak infection was noted on Day 8 with a decline by day 12. Infection was statistically significant on days 2-12. Viability of these cells was quantified by flow cytometric detection of live cells (Figure 2C). Viability in the infected condition falls to below 20% by 12 DPI. On 8 DPI, there is a statistically significant difference between viability of infected cells and uninfected cells that corresponds to the timing of peak infection. Figure 2D demonstrates representative flow cytometry plots of viability of cells over the course of infection indicating waning viability over the course of infection. Figure 2E demonstrates representative flow cytometry plots of the infection of subset of live cells on 0, 2, 5, 8, and 12 DPI.

**Figure 2.**
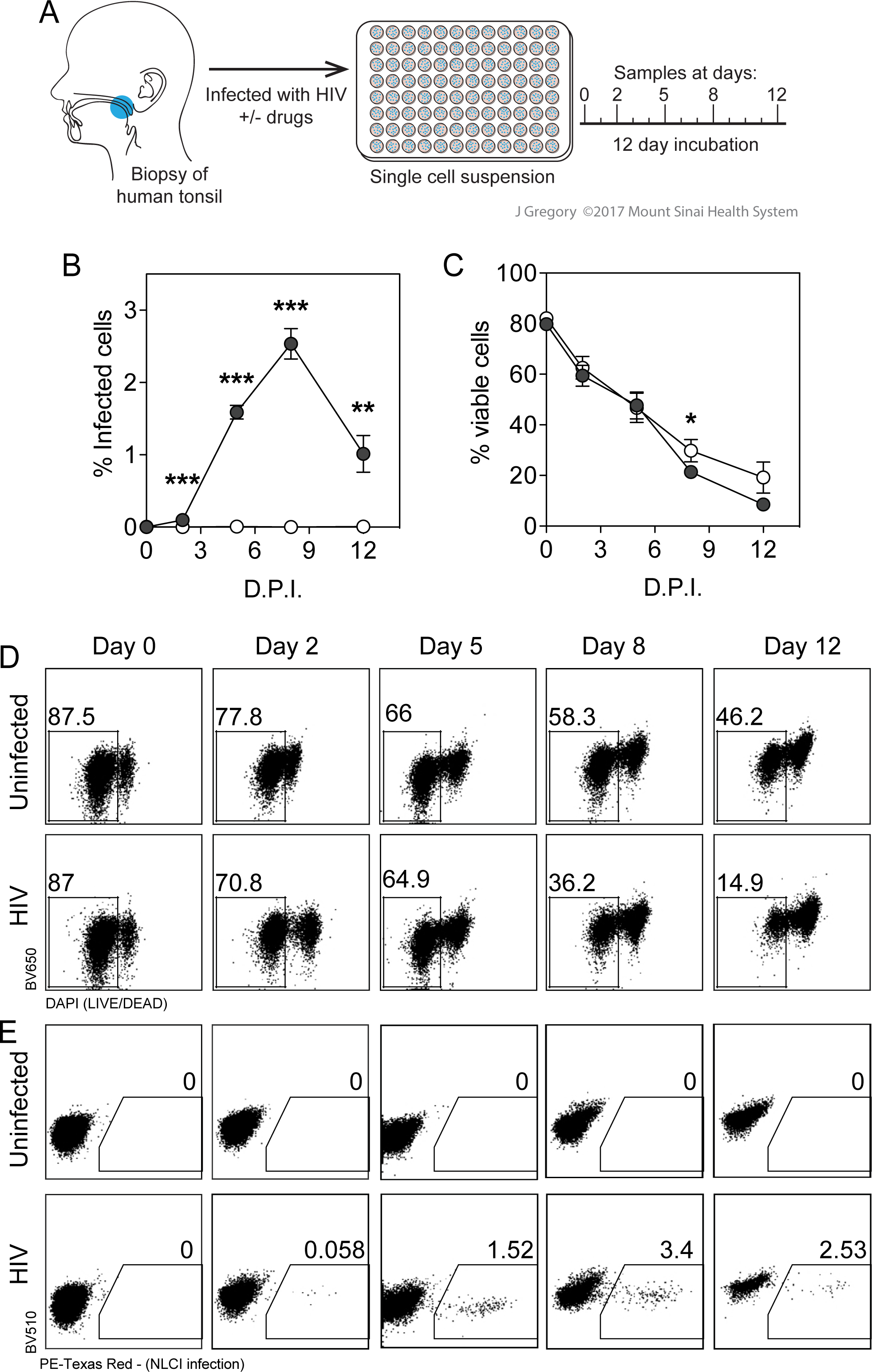
Human lymphoid aggregate culture (HLAC) of tonsil explant model supports HIV-1 infection. (A) Human tonsil explants were collected, dissected, homogenized and passed through a cell strainer. Cells were subject to Ficoll fractionation and human lymphoid aggregate cells (HLACs) were plated, then infected with HIV-1 NL-CI. Cells were collected on 0, 2, 5, 8, and 12 DPI. (B) HLACs were collected on the indicated days and analyzed by flow cytometry for productive infection by NL-CI mCherry fluorescence. Infected cells were quantified by the percentage of positive PE-Texas Red events (C) HLACs were analyzed by flow cytometry to quantify viable cells. Viable cells were quantified by the percentage of negative DAPI events (D) Representative flow cytometry plots of uninfected and infected cells are shown for LIVE/DEAD Fixable Dead Cell staining (Thermo) by flow cytometry, indicating viability of the 12 day infection time course. (D) Representative flow cytometry plots of uninfected and infected cells are shown for mCherry signal as indicative of HIV-1 productive infection. Mean values ± SEM are represented from three donors. ^∗^, p ≤ 0.05, ^∗∗^, p ≤ 0.01, ^∗∗∗^, p ≤ 0.001.

### NF449 and A438079 reduce HIV-1 productive infection in HLACs

Using this tonsil system that can support HIV-1 infection, we tested whether two P2X antagonists, NF449 (a P2X1>>P2X7 inhibitor) and A438079 (a P2X7 inhibitor) would reduce HIV-1 productive infection in comparison to AZT. HLACs were prepared as in Figure 2 and cells were harvested on 0, 2, 5, 8, and 12 DPI. Viability of these cells was quantified by flow cytometric detection of live cells (Figure 3A). As in Figure 2B, viability of cells declined over the infection course by 12 DPI. Interestingly, NF449 and AZT resulted in statistically significant increase cell survival that was most pronounced between 5-12 DPI. Figure 3B indicates infection as measured by quantification of cells with mCherry signal, as indicative of HIV-1 NL-CI productive infection. Both NF449 and A438079 at 100 μM reduce HIV-1 productive infection at all time points from 2-12 DPI comparable to inhibition seen by AZT. Surprisingly, A438079 (100 μM) effectively inhibited productive HIV-1 infection in human tonsil cells on 8 and 12 DPI, although it incompletely inhibited productive infection of PBMCs (Figure 1A). Titration of these three drugs was performed at during peak infection at 8 DPI (Figure 3C), indicating dose-dependent inhibition of HIV-1 productive infection with NF449, A438079, and AZT with IC_50_ values of 12.4 μM, 36.3 μM, and 12.0 μM, respectively.

**Figure 3.**
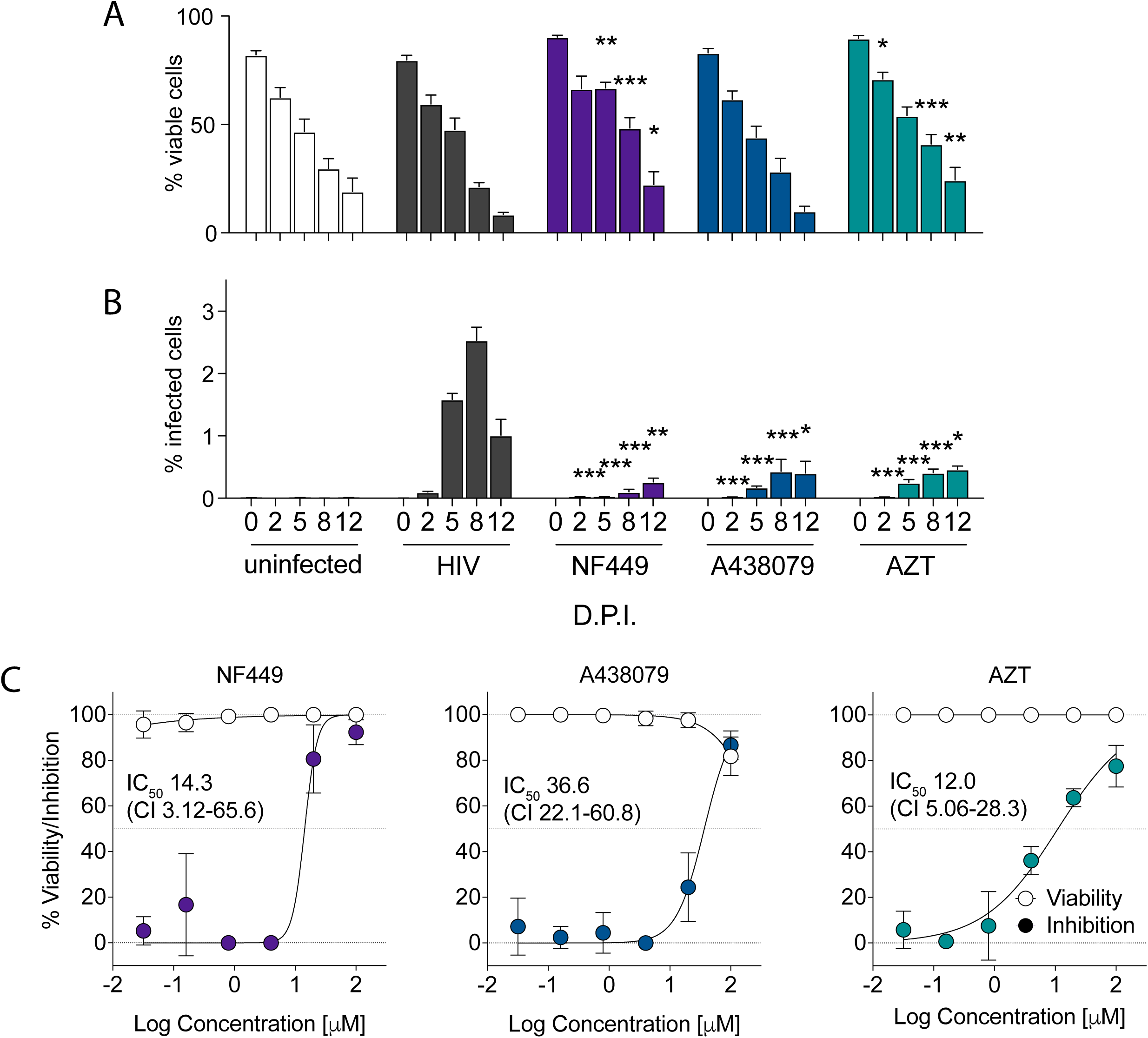
NF449, A438079, and AZT block HIV-1 replication in HLACs. HLACs were collected on 0, 2, 5, 8, and 12 DPI. (A) HLACs were analyzed by flow cytometry to quantify viable cells. (B) HLACs were analyzed by flow cytometry for productive infection by NL-CI mCherry fluorescence. (C) NF449, A438079 and AZT were tested for dose-dependent inhibition in infected HLACs at 8 DPI by a 1:5 titration down from 100 μM. Mean values ± SEM are represented from two donors ^∗^, p ≤ 0.05, ^∗∗^, p ≤ 0.01, ^∗∗∗^, p ≤ 0.001.

### HIV-1 infection is inhibited by NF449 and A438079 in human tonsil explant tissue blocks

We next tested the supernatants of human tonsil explant tissue blocks to determine how infection levels are associated with secreted cytokines. To do this, tonsils were dissected and cut into small blocks and suspended at the liquid-air interface on collagen rafts. Supernatants were collected on days 2, 5, 8, and 12 DPI (Figure 4A) and saved for experimental analysis. Media with drug were changed completely on each indicated DPI, and therefore quantification represents cumulative accumulation of viral production. While HLACs allow for more cellular analysis of viability and infection via flow cytometry, the human tonsil explant tissue block model allows for preservation of the tonsil tissue cytoarchitecture. Given prior evidence that a lymphoid tissue microenvironment is required to support the course of HIV-1 infection and pyroptosis (50, 53, 55), we pursued the development of a tonsil model that could recapitulate the observations of HIV-1 infection and stimulation of inflammatory cytokine production.

**Figure 4.**
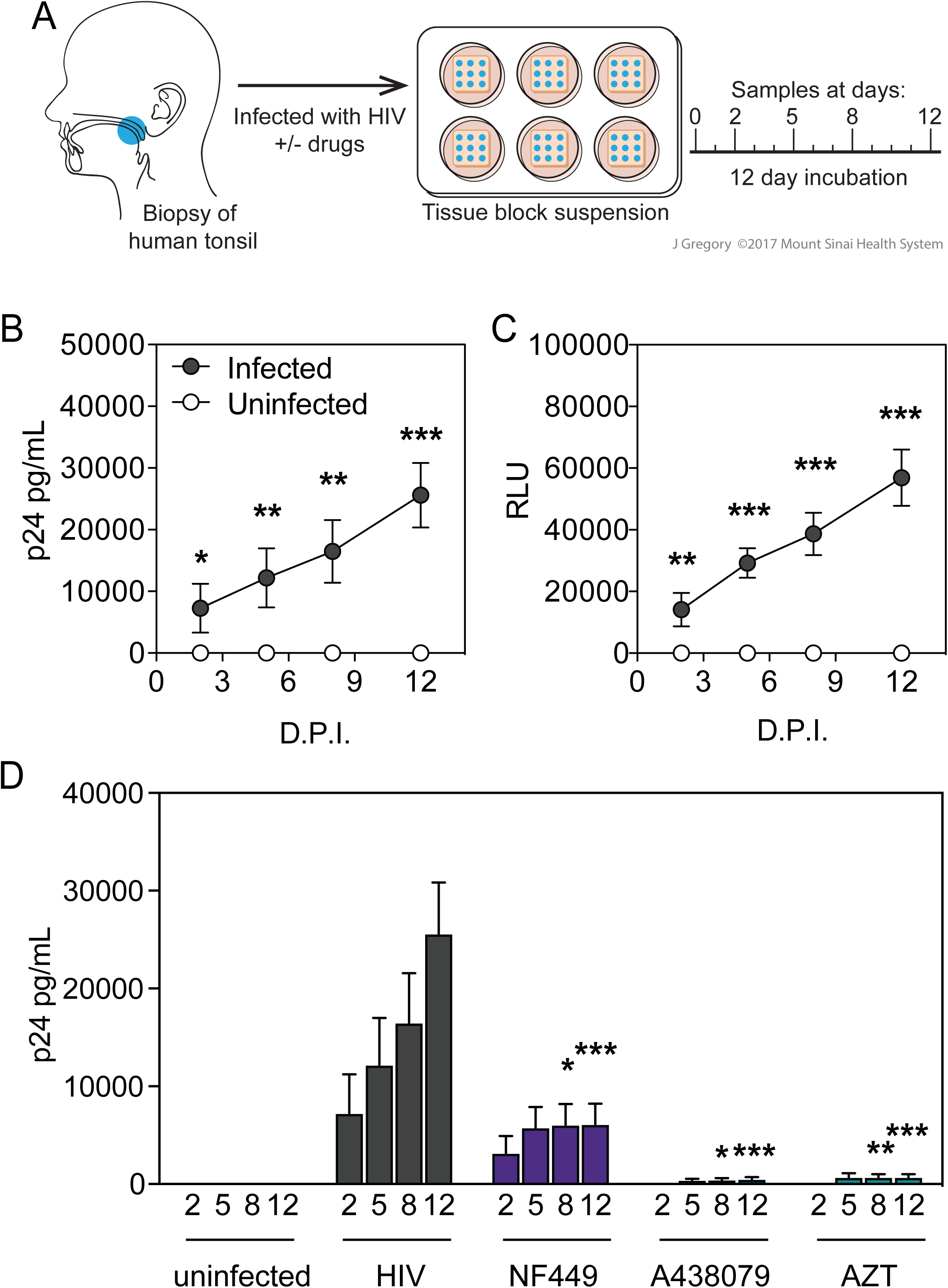
HIV-1 infection and inhibition in human tonsil explant tissue blocks. (A) Human tonsil explant tissue blocks were collected, dissected and plated in blocks on GelFoam for 24 hours prior to infection. Blocks were infected with HIV-1 NL-CI. Supernatants were collected on 2, 5, 8, and 12 DPI. (B) Supernatants collected from human tonsil explant tissue blocks at the indicated times were measured for HIV-1 p24 by ELISA. Unfilled circles indicate uninfected samples and black circles indicate infected samples. (C) Supernatants from human tonsil explant tissue blocks on each day were tested for HIV-1 infectivity as quantified by TZM-bl assay. (D) Human tonsil explants were infected with HIV-1 NL-CI in the presence or absence of indicated inhibitors (100 μM). Supernatants were collected on 2, 5, 8, and 12 DPI after infection. HIV-1 p24 antigen levels were measured by ELISA in samples in which infected tonsils were incubated in the presence or absence of inhibitors NF449, A438079, and AZT at 100μM. Data represent cumulative HIV-1 p24 production by adding the measurements at each successive time point. Mean values ± SEM are represented from six donors ^∗^, p ≤ 0.05, ^∗∗^, p ≤ 0.01, ^∗∗∗^, p ≤ 0.001.

First, we attempted to determine if HIV-1 infection and inhibition in the human tonsil explant tissue block model was comparative to the HLAC model. The HIV-1 NL-CI virus expresses an mCherry gene cloned into the *nef* position and provides and indicator of early viral gene expression. Nef expression is restored with the insertion of an internal ribosome enty site. Therefore, infection was monitored by measurement of viral antigen by HIV-1 p24 ELISA (Figure 4B) and by measuring infectivity in the supernatants by exposure to the HIV-1 indicator TZM-bl cell line and quantification of relative luminescence units (RLUs) (Figure 4C). HIV-1 infection resulted in significant p24 antigen accumulation from 2-12 DPI accompanied by infectivity of the TZM-bl cell lines.

We next tested the effect of these drugs on reducing HIV-1 p24 antigen accumulation in the *ex vivo* tonsil tissue model over the 12-day infection (Figure 4D). A significant reduction of HIV-1 p24 antigen accumulation was observed with NF449 (100 μM) treatment on 8 and 12 DPI. As in the HLAC system in Figure 3, A438079 (100 μM) inhibited HIV-1 p24 antigen accumulation on 8 and 12 DPI. In histoculture, A438079 inhibited HIV-1 productive infection to a similar extent as the positive control, AZT (100 μM) that fully blocked productive infection.

### HIV-1 infection is associated with inflammatory cytokine production in *ex vivo* lymphoid model

With confirmation of the establishment of productive infection by HIV-1 in this *ex vivo* lymphoid model, we next examined if HIV-1 infection stimulated inflammatory cytokine production. Supernatants were harvested from uninfected and infected human tonsil explant tissue blocks and were analyzed for cytokines IL-10, IL-1β, tumor necrosis factor (TNF), interleukin-12p70 (IL-12p70), interleukin-8 (IL-8), and interleukin-6 (IL-6). Supernatants were harvested over the 12-day infection time course and subject to the multiplex bead immunoassay Cytometric Bead Array (CBA; BD Biosciences) (Figure 5A). Cumulative measurements indicated that HIV-1 infection stimulated a significant increase in IL-10 and IL-1β from 2 to 12 DPI. HIV-1 infection stimulated a modest but not significant increase of TNF at 12 DPI and did not stimulate an increase in IL-12p70, IL-8, or IL-6.

**Figure 5.**
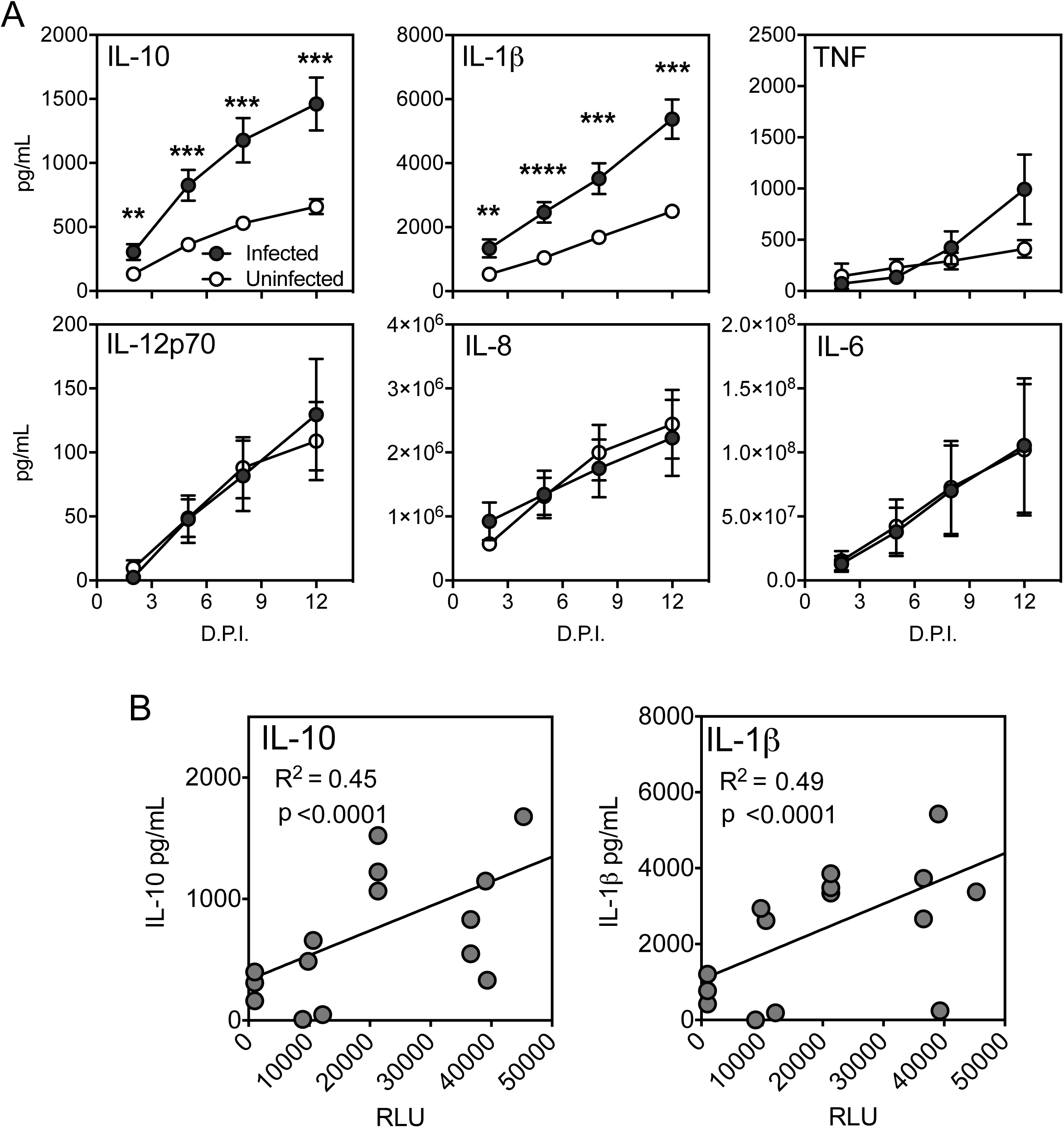
HIV-1 stimulates production of IL-10 and IL-1β in human tonsil explant model. Human tonsil explant tissue blocks were infected with HIV-1 NL-CI. Supernatants were collected on 2, 5, 8, and 12 DPI. (A) Cytokines IL-10, IL-1β, TNF, IL-12, IL-8, and IL-6 were measured in the harvested supernatants, quantified by CBA (BD Biosciences) and analyzed by flow cytometry. Data represent cumulative cytokine production by adding the measurements at each successive time point. Mean values ± SEM are represented from six donors ^∗^, p ≤ 0.05, ^∗∗^, p ≤ 0.01, ^∗∗∗^, p ≤ 0.001. (B) Quantification of TZM-bl infectivity by RLU and cytokine levels were compared by regression analysis for IL-10 and IL-1β. A positive correlation was identified between RLU and IL-10 and IL-1β. Mean values ± SEM are represented from five donors ^∗^, p ≤ 0.05, ^∗∗^, p ≤ 0.01, ^∗∗∗^, p ≤ 0.001.

We further sought to test whether the stimulation of these cytokines was associated with the magnitude of infection. Figure 5B demonstrates IL-10 and IL-1β production plotted as a function of TZM-bl infectivity. IL-10 and IL-1β production are both positively correlated with HIV-1 infection with R^2^ values of 0.45 and 0.49, respectively. This suggests that this model can support the stimulation of inflammatory cytokine production that is directly proportional to the level of HIV-1 infection. For the cytokines for which no significant difference was noted in Figure 5A, notably TNF, IL-12, IL-6, and IL-8, no correlation was observed (data not shown).

### NF449 and A438079 reduce HIV-1 stimulated IL-10 and IL-1β production in human tonsil cells

With the establishment of a tonsil system that supported HIV-1 infection and HIV-1 associated inflammatory cytokine secretion, we examined the role of purinergic signaling pathways in the induction of cytokines by HIV-1 infection. Based on our prior observations that P2X antagonists reduced HIV-1 infection and fusion (34, 57), it was of interest to test a panel of selective P2X antagonists to identify drugs with inhibition of HIV-1 infectivity in lymphocyte cell lines that would also interfere with HIV-1 stimulation of cytokines in the tonsil system. We tested whether NF449 and A438079 would reduce HIV-1-stimulated inflammatory cytokine production. Human tonsil explant tissue blocks were infected with HIV-1 NL-CI as in Figure 4. Supernatants from infected human tonsil explant tissue blocks were harvested on 2, 5, 8, and 12 DPI and analyzed for inflammatory cytokines by CBA (BD Biosciences). Cumulative IL-10 and IL-1β production over the 12-day infection course was measured in the presence or absence of indicated antagonists. HIV-1 infection stimulated IL-10 production, which was significantly reduced by NF449 and A438079 on 5, 8, and 12 DPI (Figure 6A). Additionally, HIV-1 infection stimulated IL-1β production, which was significantly reduced by A438079 at 5, 8, and 12 DPI (Figure 6B). NF449 did not significantly reduce IL-1β production until 12 DPI. AZT was not expected to inhibit either cytokine production, but did inhibit IL-10 production to a lesser extent than NF449 or A438079. AZT did not reduce IL-1β levels. These observations support the notion that P2X-selective antagonists act on HIV-associated inflammation differently than conventional ART. Overall, we conclude that NF449 and A438079 inhibit HIV-1 replication and IL-10 and IL-1β release in human tonsil explants. This suggests that P2X-selective antagonists are active in both reducing HIV-1 infection and associated inflammation.

**Figure 6.**
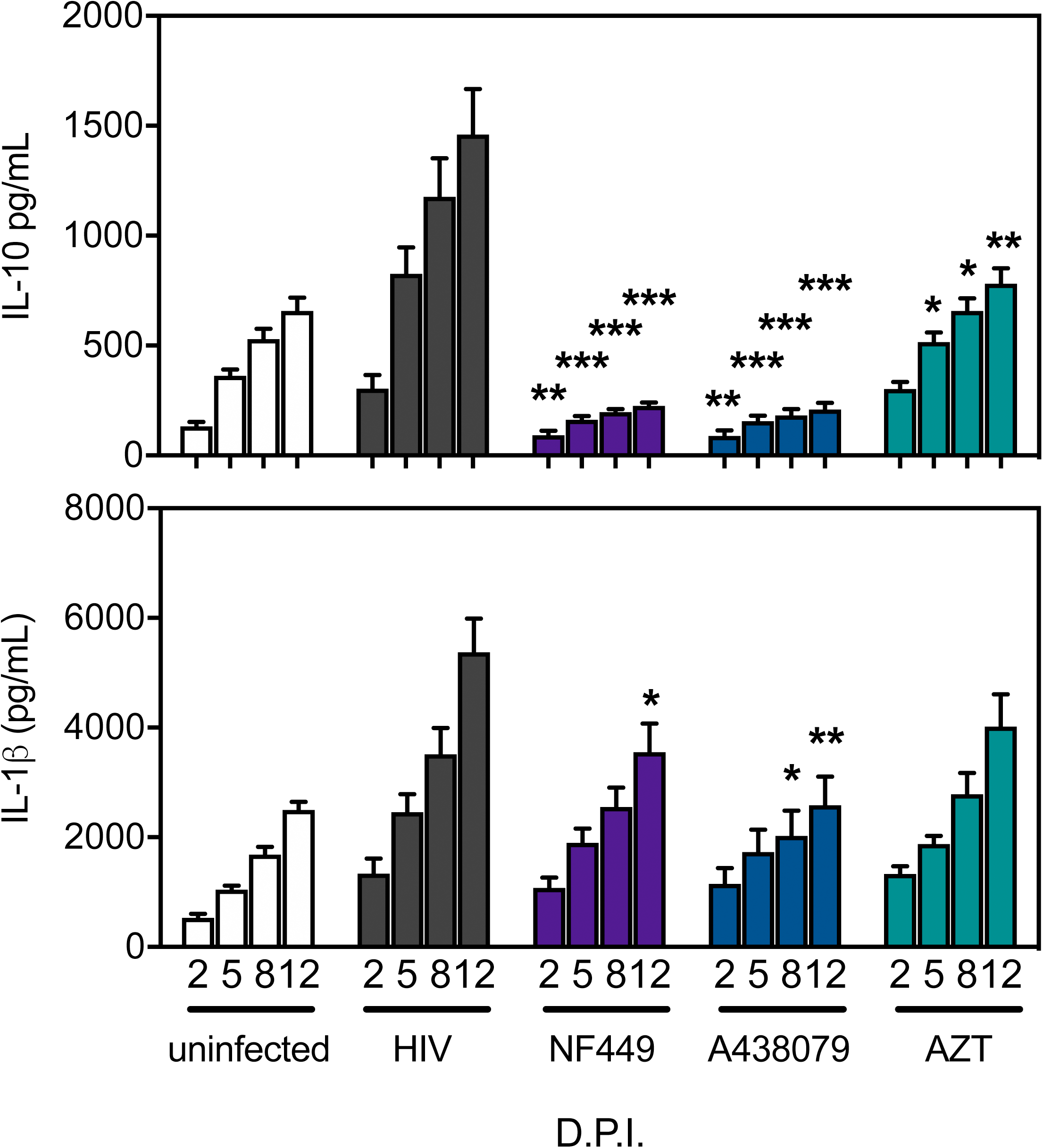
NF449 and A438079 reduce HIV-1 induced elevated cytokines IL-10 and IL-1β in *ex vivo* human tonsils. Human tonsil explants were infected with HIV-1 NL-CI in the presence or absence of indicated inhibitors (100 μM). (A) IL-10 and (B) IL-1β were measured from supernatants harvested on 2, 5, 8, and 12 DPI in the presence or absence of inhibitors NF449, A438079, and AZT. Cytokines were measured in the supernatants harvested samples and quantified by CBA (BD Biosciences) and analyzed by flow cytometry. Data represent cumulative cytokine production by summing the measurements at each successive time point. Mean values ± SEM are represented from six donors ^∗^, p ≤ 0.05, ^∗∗^, p ≤ 0.01, ^∗∗∗^, p ≤ 0.001.

## Discussion

Here we demonstrate a human *ex vivo* tonsil model that can support HIV-1 infection over a 12-day incubation. This model represents an important experimental system to test the signaling that mediates HIV-1 infection and HIV-1-stimulated inflammation and illustrates an important distinction between HIV-1 inflammation in peripheral blood and in lymphoid tissues. As soluble cytokine production is challenging to measure in PBMCs, it has been necessary for investigators to probe lymphoid tissues for evidence of HIV-1-stimulated immune activation and pyroptosis which cannot be demonstrated in peripheral blood (50–55). We demonstrate HIV-1 productive infection of human tonsils in HLACs and in human tonsil explant tissue blocks. The advantage to the HLAC system is that cell viability can be measured alongside productive infection using a fluorescent reporter virus. The human tonsil explant tissue blocks allow for measurement of soluble cytokine and the measurement of p24 antigen as an indirect measure of HIV-1 spreading infection. Using these two models, we demonstrated HIV-1 replication and HIV-1 stimulated IL-10 and IL-1β production.

In the HLAC model, we demonstrated HIV-1 productive infection occurred with a peak at 8 DPI and a corresponding decline in cell viability. All three drugs tested, NF449, A438079, and AZT reduced HIV-1 productive infection, while NF449 and AZT resulted in statistically significant increased cell survival, most notably on 8-12 DPI, which likely relates to inhibition of HIV-1 productive infection. Dose dependence inhibition of HIV-1 productive infection was noted for all three drugs with IC_50_ values all in the 10-100 μM range.

In the human tonsil explant tissue blocks, HIV-1 p24 antigen and TZM-bl infectivity steadily increased over the 12-day infection. NF449, A438079 and AZT, inhibited HIV-1 p24 and HIV-1 productive infection to statistically significant extent by 8 and 12 DPI. At those same time points, IL-10 and IL-1β cytokine stimulation steadily increased and those levels positively correlated with TZM-bl infectivity, suggesting a direct relationship between the extent of infection and IL-10 and IL-1β cytokine stimulation.

The P2X-selective antagonists tested, NF449, and A438079, reduced HIV-1-stimulated levels of IL-10 with statistical significance between 5-12 DPI. Treatment with AZT at the corresponding time points resulted in less inhibition of IL-10 compared to the infected condition. This suggests that inflammatory signaling may be blocked not directly through inhibition of productive infection, but through alternative signaling mechanisms. These observations highlight the unique properties of NF449 and A438079 as novel agents that reduce inflammatory changes independent of their antiviral properties.

Additionally, the drugs were tested for effect on HIV-stimulated IL-1β production. NF449 reduced HIV-1-stimulated levels modestly at 12 DPI whereas A438079 reduced HIV-1-stimulated levels of IL-1β at 8-12 DPI. AZT did not reduce IL-1β levels. Of note, A438079 did not have full inhibition of HIV-1 productive infection in PBMCs, but unexpectedly had strong inhibition of HIV-1 p24 accumulation in human tonsil explants and reduced IL-10 and IL-1β secretion. This surprising observation suggests that the nature of A438079 inhibition of productive infection in tonsils may not be ascribed to a direct link between A438079 and HIV-1 entry, but rather through cytokine-dependent signaling. There are several possible explanations for this phenomenon. The immunomodulatory role of IL-10 may serve to enhance HIV-1 permissively, but when levels of IL-10 are reduced by A438079, cells may be more susceptible to A438079 inhibition of HIV-1 infection. Tonsil tissue represents mixed cellular populations with stromal compartments with altered sensitivity to P2X inhibition as compared to PBMCs (58–60). It will be of interest to explore the cell-type heterogeneity of HIV-1 infection in tonsils to determine the cell type and signaling mechanisms that drive IL-10 and IL-1β production. These directions may lead to the development novel therapeutic agents that retain inhibition of spreading HIV-1 infection and can reduce HIV-1 stimulated levels of IL-10 and IL-1β.

The role of P2X receptors in HIV-1 pathogenesis likely relate to downstream activation of the NLRP3 inflammasome. The NLRP3 activates immature caspase-1 to activated caspase-1, which cleaves and activates pro-IL-1β. These cytokines, among others, play a pivotal role in signaling of other inflammatory cytokines. Emerging evidence suggests a key role for the inflammasome in atherosclerotic disease progression (61–63) (64–71) and in HIV-1 disease (46, 72–77). Inflammasome activation requires two signals, one for priming (i.e. TLR signaling which results in transcriptional regulation), and then activation for inflammasome complex assembly. Together with TLR signaling, P2X7 can signal inflammasome activation and subsequent IL-1β release (78). As the tonsil tissue contains soluble factors that likely include TLR agonists, inflammasome activation is readily measurable with second signal stimulation, i.e. P2X7 activation by HIV-1 infection. HIV-1 infected patients have elevated circulating levels of LPS, which can serve to increase transcriptional activation of pro-inflammatory cytokines (11,49). Elevated levels of IL-1β are associated with many of the AANCCs seen in HIV-1 infected patients (79–82). Currently, there are multiple clinical trials assessing the safety and efficacy of anti-IL-1β antibodies in cardiovascular disease (83, 84). Canakinumab, a human monoclonal IL-1β antibody, has been shown to significantly decrease arterial inflammation in HIV-1 infected individuals (85). The role of P2X receptors in the secretion of IL-1β may represent a key mechanism for HIV-1 associated inflammation. Elevated IL-1β is observed in HIV-1 infected patients (76, 86–88) and an emerging literature implicates the role of IL-1β in atherosclerotic cardiovascular disease in both HIV-1 infected and uninfected patients (83–85). Intriguing studies in CD4^+^ T cells find that pathogen sensor interferon-γ-inducible protein 16 (IFI-16) recognition of HIV-1 DNA can activate the NLRP3 inflammasome that induces pyroptosis and may represent a mechanism for CD4^+^ T cell depletion in HIV-1 disease and progression to advanced disease (53–55).

The fact that both IL-10 and IL-1β were together stimulated by HIV-1 infection and reduced by NF449 and A438079 are surprising, given that they have opposing inflammatory effects. IL-10 is an immunomodulatory cytokine with immunosuppressive activities and has been implicated in immune exhaustion and cell death through inhibition of NF-kB activity (89–93). IL-10 activation has been linked to P2X7 signaling with the observation of down-modulation of IL-10 receptor expression with P2X7 activation (94). IL-10 has the potential to impact many areas of HIV-1 infection including CD4 function, chemokine receptor expression, and modulation of replication (95–97). IL-10 gene polymorphisms and epigenetics have been shown to be associated with variations in HIV-1 transmission and disease progression, as long-term non-progressors (LTNP) have low IL-10 levels compared with HIV-1 progressors (98–103). This suggests that IL-10 may be important for modulating the immune response to HIV-1 infection and the high levels of HIV-1-stimulation may account for the relatively low levels of induction of other pro-inflammatory cytokines such as IL-6 and TNF. Further studies are needed to understand the role of IL-10 in HIV-1 disease progression and inflammation, and P2X-selective antagonists may play a role in developing novel therapeutics that target both HIV-1 infection and inhibition of IL-10 signaling.

Figure 7A summarizes the observations of HIV-1 infection and inflammatory cytokine production in both PMBCs and tonsils. While all drugs inhibit HIV-1 infection, A438079 had the least effect on inhibition of HIV-1 infection. It should be noted that IL-10 and IL-1β stimulation were not observed in this system. Since it was not possible to demonstrate HIV-1 specific stimulation of inflammatory cytokines in this model, it was necessary to establish a lymphoid model that would more accurately recapitulate inflammatory cytokine signaling. In the tonsil model, all three drugs inhibited HIV-1 infection to a comparable magnitude in HLACs, while A438079 and AZT inhibited p24 accumulation more than NF449. By comparison, NF449 and A438079 inhibited IL-10 production most strongly while AZT only modestly reduced IL-10 production. Finally, NF449 and A438079 inhibited IL-1β production modestly while AZT did not inhibit IL-1β production.

**Figure 7.**
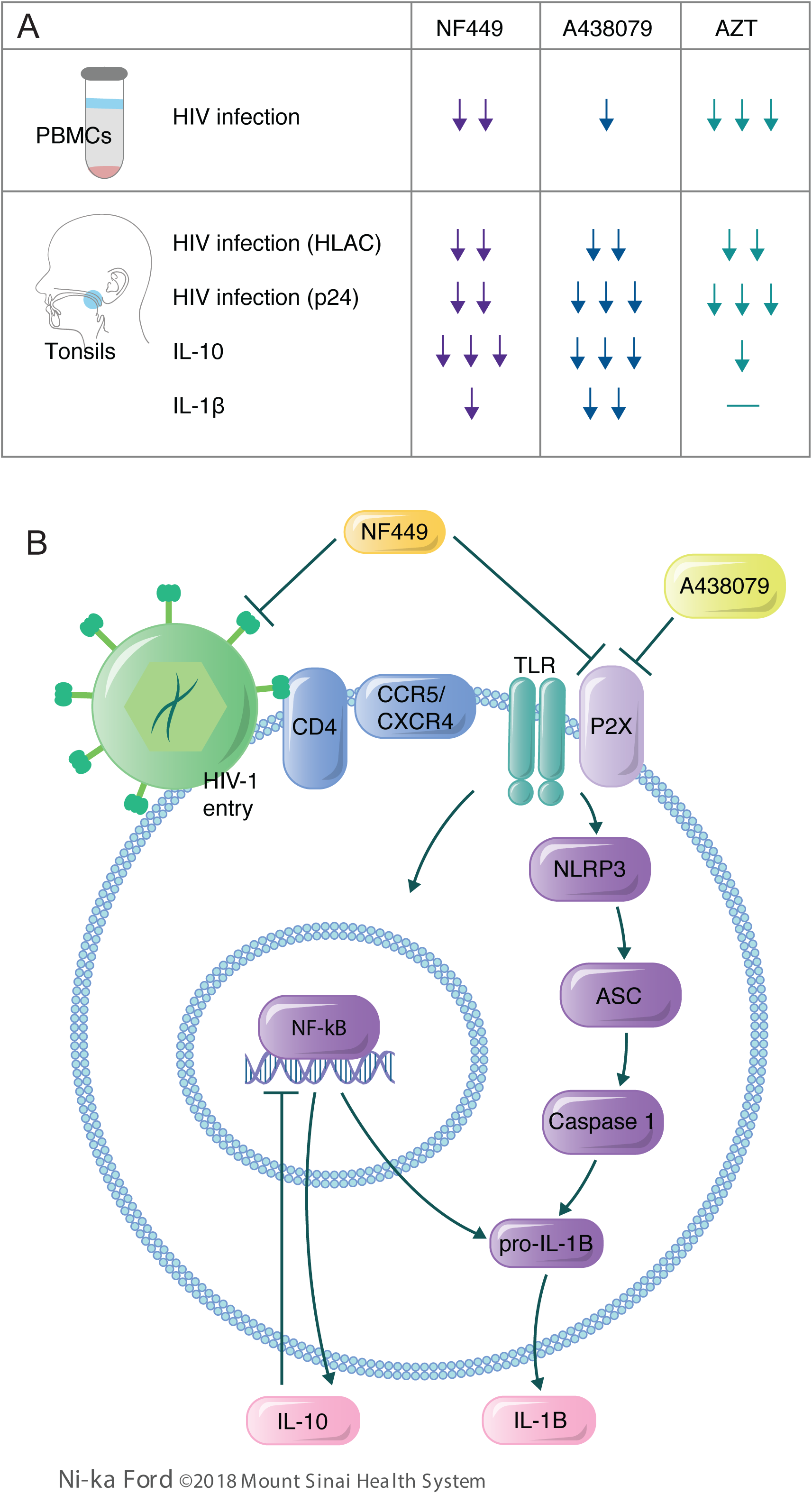
Model for signaling in HIV-1 mediated inflammation. (A) Table summarizing observations from drug activity across PMBC and tonsil models. Arrows indicate relative magnitude of effect of inhibition in each of the models depicted. (B) NF449 inhibits HIV-1 productive infection and downstream signaling of the NLRP3 inflammasome. This pathway is activated in concert with TLR4 receptor activation which can drive caspase-1 to cleave pro-IL-1β to mature IL-1β. Mature IL-1β can be secreted or induce pyroptosis in CD4^+^ T-cells. This signaling can also activate NF-kB dependent transcriptional regulation of IL-10. Inhibition of this mechanism by NF449 may explain enhanced cell survival in tonsil cells. A438079, by contrast, may act directly on P2X7 to inhibit receptor signaling that is required for HIV-1 entry. P2X7 inhibition results in the inability to activate the NLRP3 inflammasome and pyroptosis in tonsil cells, which does not occur in PBMCs. Therefore, A438079 depends on intact inflammasome signaling to exert inhibition on HIV-1 productive infection.

Taken together, we propose the model in Figure 7B, in which P2X-selective antagonists play a role in inhibiting both HIV-1 infection and HIV-1-stimulated inflammation. The model indicates that NF449 and A438079 may have different mechanisms of action on HIV-1 entry and inflammation. Stimulation of inflammatory signaling by P2X7 and TLR4 results in NLRP3-dependent production of IL-1β as well as NF-kB-dependent regulation of IL-10. NF449 inhibits this inflammation and HIV-1 productive infection in both PBMCs and tonsil cells, suggesting that the inhibition is not dependent on intact inflammasome signaling, but may act more directly on HIV-1 entry mechanisms. By contrast, A438079 inhibits HIV-1 productive infection that is limited to the tonsil system, indicating that this inhibition is dependent on intact NLRP3 signaling mechanisms that are not activated in PBMCs. The role of cytokine signaling in permissivity to HIV-1 infection in mixed cell populations is an important area of future investigation as the tonsil tissue model system is important in understanding the interplay between HIV-1 pathogenesis and the immune response.

Here we demonstrate that P2X-selective antagonists have the potential to reduce HIV-1 infection and HIV-1-stimulated inflammatory cytokine production. We conclude from these studies that P2X-selective antagonists may represent potential HIV-1 therapeutic options that serve to inhibit HIV-1 replication and innate immune sensing. Further studies will be necessary to identify selective inhibitors that are amenable to pharmacologic development and the precise mechanism of their inhibition, but these observations introduce important prospects for dually active therapeutic options that would reduce the burden of morbidity and mortality of chronic inflammation in HIV-1-infected individuals.

## Materials and Methods

### Virus production

HIV-1 NL-CI contains mCherry in place of *nef*, and *nef* expression is directed by a downstream internal ribosome entry site (IRES) (44). Pseudoviruses were produced by co-transfecting 293T/17 cells with HIV-1 *rev*- and env-expressing plasmids and the pNL4-3Δenv R-E-plasmid using the jetPEI transfection reagent (Polyplus-transfect SA). Supernatants were harvested after 48 hours and clarified by high-speed centrifugation (Sorvall ST 40R Centrifuge, Thermo Fisher Scientific) at 100,000 × g at 4°C for 2 hours and 0.45 μm filtration. Single-use aliquots were stored at –80°C. Viral stocks were quantified via enzyme-linked immunosorbent assay (ELISA), as described below. HIV-1_MN_ (X4-tropic) was produced at the AIDS Vaccine Program, National Cancer Institute as previously described (104–106).

### Cells and cell lines

PBMCs were obtained from de-identified HIV-1 negative blood donors (New York Blood Center), purified by Ficoll (HyClone) density gradient centrifugation and were maintained in RPMI 1640 medium (Sigma) containing 10% fetal bovine serum (FBS; Sigma), 100 U/ml penicillin (Gibco), 10 U/ml streptomycin (Gibco), and 2 mM glutamine (Gibco) (complete RPMI). The 293T/17 cell line was used to produce pseudoviruses and purchased from the American Type Culture Collection. The TZM-bl cell line was obtained from Dr. John C. Kappes, Dr. Xiaoyun Wu, and Tranzyme Inc. through the NIH ARP. The 293T/17 and TZM-bl cells were maintained in Dulbecco’s modified Eagle Medium (DMEM; Sigma) containing 10% cosmic calf serum (CCS; HyClone) and 100 U/mL penicillin, and 10 U/mL streptomycin, and 2 mM glutamine (Gibco) (complete DMEM).

### Establishment of human explant tonsil model and processing of HLACs

Human tonsils were collected from routine tonsillectomy performed at the Mount Sinai Health System in New York by BT under an Institutional Review Board-approved protocol. Tonsils were collected within several hours of surgery and dissected into 2 mm tissue blocks. Human tonsil explant tissue blocks were plated 9 per well atop a collagen sponge GelFoam (Pfizer) in a 6-well plate in ~3 mL media (Costar), as previously described (56). Media was completely changed every 2-3 days with or without indicated inhibitor and saved in aliquots at –80°C for further experiments. For HLAC experiments, dissected tissue was passed through a 40-μm cell strainer and purified by Ficoll density-gradient centrifugation, as previously described (51). Cells were plated at 2.5 × 10^5^ cells/well in a round-bottom 96-well plate (Costar) and spun down at 500 × g for 3 minutes every 2-3 days to replace media with our without indicated inhibitor. Human tonsil explant tissue blocks and HLACs were maintained in RPMI 1640 medium (Life Technologies) containing 15% FBS, 2 mM GlutaMAX (Life Technologies), 2 mM L-glutamine (Corning), 1 mM sodium pyruvate (Corning), 1% MEM non-essential amino acids (Corning), 2.5 μg/mL Amphotericin B (HyClone), 50 mg/mL gentamicin sulfate (Corning), and 0.3 mg/ml Timentin (bioWORLD).

### Antagonists

Inhibitors were tested for the ability to block HIV-1 infection and HIV-1 associated inflammation at 100 μM, unless otherwise stated. These include NF449 (Tocris), a P2X1 selective antagonist, A438079 (Tocris), a P2X7 selective antagonist, and the reverse transcriptase inhibitor azidothioidine (AZT; Sigma). NF449 and A438079 were diluted from 10 mM stocks reconstituted in DMSO while AZT was diluted from 10 mM stocks reconstituted in water.

### Flow cytometry and gating strategy

An LSR II flow cytometer (BD Biosciences) was used to detect infection and viability in PBMCs and HLACs. Viable cells were detected with LIVE/DEAD Fixable Dead Cell Stain (Life Technologies), an amine reactive fluorescent dye that can penetrate the membranes of dead cells but not live cell membranes. Samples were stained with LIVE/DEAD Fixable Blue Dead Cell Stain or LIVE/DEAD Fixable Violet Dead Cell Stain at a concentration of 1:1000 in Wash Buffer (PBS supplemented with 2 mM EDTA and 0.5% bovine serum albumin). Stained cells incubated at 4°C for 30 minutes, then were washed and fixed in 2% paraformaldehyde for flow cytometry. All cells were initially discriminated by side scatter (SSC) area versus forward scatter (FSC) area (SSC-A/FSC-A); doublets were excluded using FSC height (FSC-H) vs FSC-A. Viability was determined by gating on negative populations for LIVE/DEAD Fixable Dead Cell Stain. Infection was detected by the presence of mCherry in cells infected with HIV-1 NL-CI. mCherry was detected using the phycoerythrin-Texas Red (PE-Texas Red) channel, LIVE/DEAD Fixable Violet Dead Cell Stain was detected with the 3-carboxy-6,8-difluoro-7-hydroxycoumarin (Pacific Blue) channel, and LIVE/DEAD Fixable Blue Dead Cell Stain was detected with the 4’,6-diamidino-2-phenylindole (DAPI) channel. All cells within a single experiment were analyzed using the same instrument settings. Flow cytometry data were exported and analyzed using FlowJo software, version 9.3.2 (Tree Star, Ashland, OR).

### Productive infection in PBMCs

PBMCs were activated with PHA (4 μg/ml) and IL-2 (50 U/mL) for 3 days and infected by spinoculation, as previously described (107, 108). Briefly, 2.5 × 10^5^ cells were incubated in the presence or absence of indicated inhibitors in a 96-well flat bottom plate for 30 minutes at 37°C then spun at 1,200 × *g* for 99 minutes with 47.7 ng HIV-1 NL-CI. After overnight incubation at 37°C, the culture medium was replaced with complete RPMI containing IL-2 (50 U/ml) and 10 μM AZT. At 48 h after spinoculation, cells were stained and fixed in 2% paraformaldehyde for flow cytometry, as described above.

### *Ex vivo* infection of human tonsil explant tissue blocks and HLACs

Human tonsil explant tissue blocks from each donor were individually inoculated with 5 μl of HIV-1 NL-CI (equivalent to 3.24 ng p24),or left uninfected in the presence or absence of indicated inhibitors. We measured HIV-1 p24 antigen in harvested supernatants and infectivity in relative RLUs by TZM-bl assay as described below, as the NL-CI fluorophore can only be detected in cell-based systems. Data are expressed as a cumulative value to account for total successive media changes. For HLAC experiments, cells were incubated in the presence or absence of indicated inhibitors for 30 minutes at 37°C before infection with 25 ng HIV-1 NL-CI p24 per well. Cells were stained and fixed for flow cytometry, as described above.

### p24 ELISA

Viral stocks and tonsil tissue supernatants were quantified via ELISA with coating antibody D7320, sheep anti-HIV-1-p24 gag (Aalto Bio Reagents), as previously described (57, 109). Briefly, anti-p24 capture antibody was coated on high binding plates (Costar) at 1:200 in 0.1 M NaHCO_3_. After overnight incubation at room temperature, plates were blocked with 2% nonfat dry milk for 1 h. HIV-1 samples were treated with 1% Empigen and added to wells, along with titration of p24 standard, at room temperature for 3 h. Alkaline phosphatase conjugated mouse anti-HIV-1 p24 (CLINIQA) was added (1:8000 in TBST 20% sheep serum) and incubated for 1 h. Plates were developed with Sapphire Substrate (Tropix) and measured on Fluo Star Optima plate reader.

### TZM-bl HIV-1 infectivity assay

Virus infectivity of supernatants collected from tonsil was measured using a β-galactosidase-based luciferase assay (Promega) with TZM-bl target cells, as previously described (110). Briefly, TZM-bl cells were plated at 1.5 × 10^4^ cells/well in a flat-bottom 96-well plate (Costar). Harvested supernatants (containing 0.1 ng HIV-1 p24) were added to each well then incubated at 37°C. Media was exchanged 24 h after incubation and a luciferin-galactoside substrate (6-O-β-galactopyranosyl-luciferin) was added after 48 h. The cleavage of the substrate by β-galactosidase generates luminescent signals measured in RLUs. Each test and control condition was tested in duplicate or triplicate. Assay controls included replicate wells of TZM-bl cells alone (cell control). The virus inputs were the diluted virus stocks yielding equivalent RLUs (typically ~100,000 RLUs) under the different assay conditions. The RLU present in uninfected samples were subtracted as background for all samples for each time point.

### Cytokine measurements

PBMCs were isolated from patient samples and 2.5 × 10^5^ cells per well in a 96 well plate were incubated in the presence or absence of dilutions of inhibitor for 30 minutes. HIV-1_MN_ (X4-tropic) was added at 300 ng/ml in the presence or absence of 1 pg/ml LPS (Sigma) and incubated for 12 hours. Supernatants were collected from PBMCs or samples of tonsil tissue and were analyzed for IL-10, IL-1β, TNF, IL-12p70, IL-8, and IL-6 using BD^™^ CBA Human Inflammatory Cytokines Kit (BD Biosciences). Standard curves were generated and cytokine concentrations were extrapolated using the FCAP Array software (BD Biosciences). The measurements indicated are representative of 3 separate PBMC donors and 6 separate tonsil donors.

### Statistical analysis and calculations

Comparisons were performed using GraphPad Prism 7, version 7.0d (GraphPad Software). DMSO-treated controls were set to 100% and drug-treated conditions were expressed as a percentage of control. Statistical analyses were performed on inhibition data that reached ≥50% with a one-tailed student’s t-test. A *P* value of less than 0.05 was considered statistically significant.

### Study Approval

ISMMS IRB protocol number 06-0980 was approved by the Program for the Protection of Human Subjects Institutional Review Board of the Mount Sinai Health System (New York, New York, USA). All patients participating in tonsil analysis gave written informed consent.

## Author contributions

AYS and NDD designed and performed experiments. AYS and NDD carried out productive infection assays, RG, MO, and NB provided reagents and guidance on cytokine measurements. BT contributed the tonsils and KWH, JAB, and JKL assisted AYS in protocol development and tonsil processing. AYS and THS wrote the paper. THS and BKC conceived the approach.

## Acknowledgements

We would like to thank the members of the Chen laboratory for meaningful discussions. Many thanks to Maxwell Allison and Elizabeth Osota for their help in preparing samples for tonsil tissue. T. Swartz was funded by the NIH K08AI20806 and by the Schneider-Lesser Foundation. This work was supported by grants to B. Chen from NIH, NIAID R01AI074420, NIDA Avant Garde DP1DA028866.

## Notes

The authors have declared that no conflict of interest exists.

